# Maintenance of immune tolerance during aging requires competitive expansion of memory phenotype regulatory T cells

**DOI:** 10.64898/2026.06.16.732599

**Authors:** Bo Chin Chiu

## Abstract

Memory phenotype (MP) CD4^+^ T cells accumulate with age; however, the immunological significance of this population shift and the mechanisms that preserve MP T-cell fitness remain poorly understood. Here, we show that age-associated accumulation of MP T cells is regulated by intra-clonal competition and that competitive expansion of MP regulatory T (Treg) cells is required for maintenance of immune tolerance during aging. Using MLL1 deficiency as a model, we found that Mll1-deficient T cells failed to undergo normal age-associated accumulation and were progressively outcompeted by wild-type (WT) cells despite retaining the ability to proliferate and generate MP populations in the absence of competitors. Mechanistically, MLL1 preserved competitive fitness by maintaining transcription of TCRα variable region genes and sustaining T-cell receptor expression during proliferation. Loss of competitive fitness impaired MP Treg-cell expansion and disrupted immune homeostasis. Remarkably, a small population of WT Treg cells restored tolerance through extensive expansion of MP Treg cells. These findings identify competitive expansion of MP Treg cells as a critical mechanism for maintaining immune tolerance during aging.

## Introduction

Memory phenotype (MP) CD4^+^ T cells are a population of steady-state T cells that lack expression of lymph node homing receptors such as CD62L and CCR7 and are found in both humans and animal models [1-3]. A defining feature of MP T cells is their high rate of turnover [2-4]. MP cells are present within both conventional CD4^+^ T cells and regulatory T (Treg) cells, with MP Treg cells exhibiting substantially higher turnover rates than MP conventional CD4^+^ T cells. Studies in mice have shown that MP T cells arise through spontaneous proliferation driven by self and environmental antigens [1, 5-8]. Importantly, spontaneous proliferation is distinct from homeostatic proliferation. Whereas spontaneous proliferation is driven primarily by antigen recognition and co-stimulation, homeostatic proliferation is driven largely by common γ-chain cytokines under lymphopenic conditions [6-8].

Spontaneous proliferation and MP-cell development are tightly regulated by competition among T cells for limiting stimulatory signals [1, 6]. Because this competition is mediated by the T cell receptor (TCR), it occurs primarily among T cells sharing similar antigen specificities and is therefore referred to as intra-clonal competition. During thymic development, intra-clonal competition plays a central role in establishing the balance between naïve Treg cells and autoreactive conventional CD4^+^ T cells [9, 10]. Whether similar competitive mechanisms contribute to maintenance of immune tolerance within the peripheral MP compartment remains unclear.

MP T cells accumulate progressively with age, whereas naïve T cells decline as thymic output decreases [11-13]. Consequently, maintenance of the peripheral CD4^+^ T-cell compartment becomes increasingly dependent on MP T cells. This shift is particularly pronounced within the Treg compartment, where MP Treg cells expand substantially with age [14, 15]. Because immune tolerance depends on the balance between autoreactive conventional CD4^+^ T cells and Treg cells [16], preserving this balance over time relies on the fitness and expansion of MP populations. Whether age-associated expansion of MP Treg cells merely reflects immune aging or instead represents an essential mechanism for maintaining immune tolerance remains unknown. Likewise, the contribution of intra-clonal competition to maintenance of tolerance in aged animals has not been defined.

MLL1 is a ubiquitously expressed epigenetic regulator that functions as a mitotic bookmarking protein [17-19]. By remaining associated with actively transcribed genes during cell division, MLL1 facilitates restoration of transcription in daughter cells. Previous studies have demonstrated important roles for MLL1 in multiple CD4^+^ T-cell subsets, including Th1, Th2, and Treg cells [18–20]. We therefore used MLL1 deficiency as a model to investigate how competitive fitness of MP T cells is maintained during aging. Here, we show that MLL1 preserves the competitive fitness of MP CD4^+^ T cells by maintaining T-cell receptor (TCR) expression during proliferation. Although Mll1-deficient T cells retain the capacity to proliferate and generate MP populations, they are progressively outcompeted by wild-type cells because of impaired maintenance of TCR expression. This defect is particularly pronounced in Treg cells, where loss of competitive fitness prevents normal expansion of the MP compartment and disrupts immune homeostasis. Our findings identify competitive expansion of MP Treg cells as a critical mechanism for maintaining immune tolerance during aging and reveal how epigenetic maintenance of TCR expression supports this process.

## Results

### MLL1 promotes the accumulation of memory phenotype (MP) CD4^+^ T cells with age

To investigate the role of MLL1 in peripheral T cells, we generated T cell-specific MLL1-deficient (Mll1KO) mice by crossing mice carrying a floxed Mll1 allele with CD4-Cre transgenic mice. Consistent with previous studies, MLL1 deficiency had minimal effects on thymic T cell development and peripheral T cell maintenance [20]. Because MLL1 regulates the function of multiple CD4^+^ T cell subsets, including Th1, Th2, and Treg cells [21-23], alterations in the immune environment of Mll1KO mice could indirectly influence T cell homeostasis. To distinguish cell-intrinsic effects from environmental influences, we generated mixed bone marrow chimeras using congenically marked Mll1KO (CD45.2^+^) and wild-type (WT; CD45.1^+^) donor cells. Mixed bone marrow cells were transferred into lethally irradiated CD45.1^+^CD45.2^+^ WT recipients, and donor-derived T cells were analyzed by flow cytometry following reconstitution (Figure 1A). Congenic markers enabled discrimination of residual host cells (CD45.1^+^CD45.2^+^), WT donor cells (CD45.1^+^), and Mll1KO donor cells (CD45.2^+^) (Figure 1B). Three months after reconstitution, the frequency of memory phenotype (MP) CD4^+^ T cells, identified by loss of CD62L expression, was reduced among Mll1KO donor cells compared with WT donor cells in both peripheral lymph nodes (PLN; Figure 1C) and spleen (Figure 1D). These findings suggest that MLL1 promotes the accumulation of MP CD4^+^ T cells.

**Figure 1.**
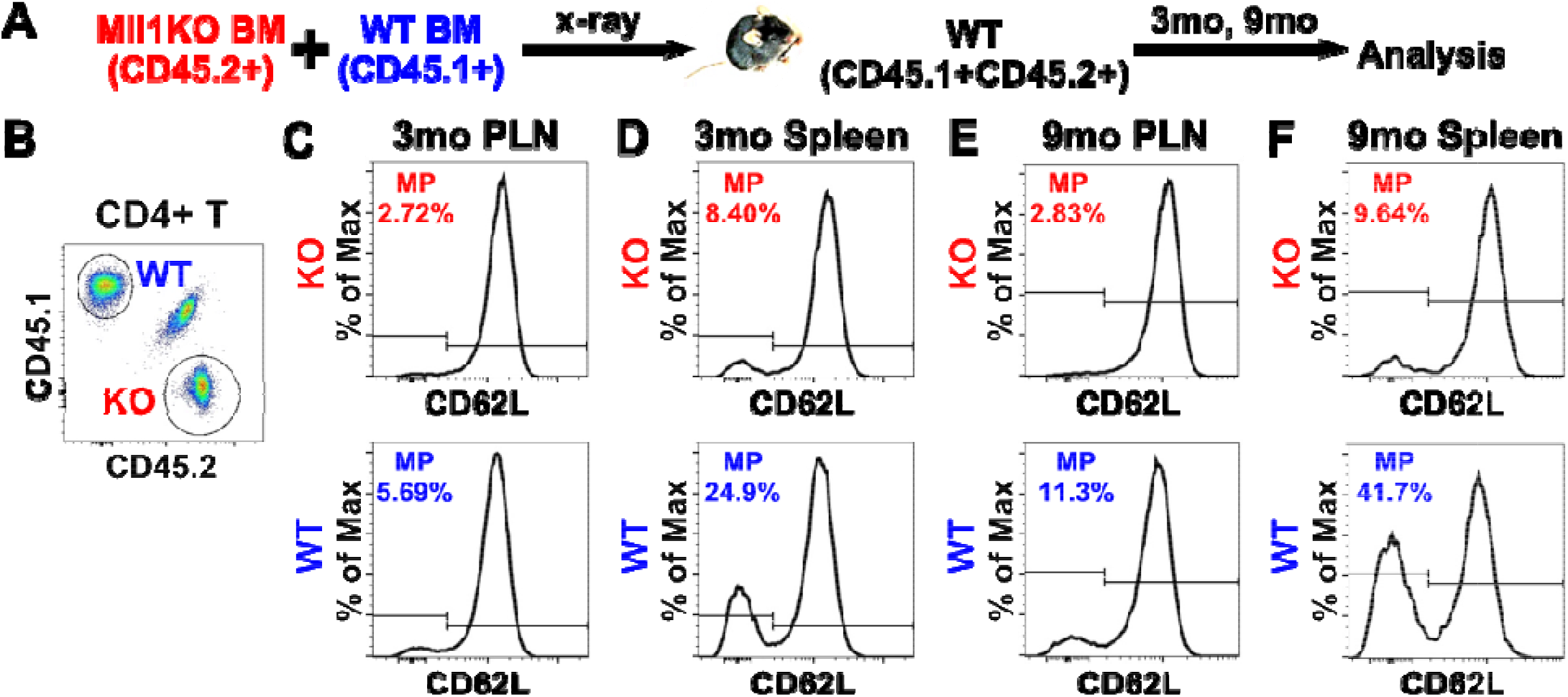
MLL1 promotes the accumulation of memory phenotype (MP) CD4^+^ T cells with age. Bone marrow cells from congenically marked Mll1KO (CD45.2^+^) and wild-type (WT; CD45.1^+^) mice were mixed and transferred into lethally irradiated WT (CD45.1^+^CD45.2^+^) recipients. Donor-derived CD4^+^ T cells were analyzed by flow cytometry 3 months (C, D) or 9 months (E, F) after reconstitution. (A) Experimental design. (B) Representative flow cytometric analysis of CD4^+^ T cells from peripheral lymph nodes showing CD45.1 and CD45.2 expression. Mll1KO (CD45.2^+^) and WT (CD45.1^+^) donor cells are distinguished by congenic markers. (C-F) CD4^+^ T cells from peripheral lymph nodes (C, E) and spleen (D, F) were analyzed for CD62L expression. MP cells were defined by loss of CD62L expression. Panels C and D show analysis at 3 months, and panels E and F at 9 months after reconstitution. Data are representative of three independent experiments.

Thymic involution results in a progressive decline in naïve T cell production with age, while the peripheral MP T-cell compartment gradually expands [11-13]. To determine whether MLL1 contributes to this age-associated accumulation, mixed chimeras were maintained for an additional six months before analysis. Strikingly, the frequency of MP CD4^+^ T cells among Mll1KO donor cells remained low in both PLN (Figure 1E) and spleen (Figure 1F), whereas WT donor cells exhibited a marked increase in MP T-cell frequency over time. Consequently, the difference between WT and Mll1KO cells became substantially more pronounced in older chimeras. These findings indicate that MLL1 is required for the normal age-associated accumulation of MP CD4^+^ T cells.

### MLL1 promotes participation in spontaneous proliferation under competitive conditions

Memory phenotype (MP) CD4^+^ T cells arise through spontaneous proliferation under steady-state conditions [6]. To determine whether MLL1 contributes to this process, congenically marked naïve Mll1KO (CD45.2^+^) and wild-type (WT; CD45.1^+^) CD4^+^ T cells were mixed, labeled with CFSE, and co-transferred into WT (CD45.1^+^CD45.2^+^) recipients (Figure 2A). Five weeks later, donor-derived cells were analyzed by flow cytometry. Congenic markers enabled discrimination of Mll1KO, WT, and host-derived T cells (Figure 2B). The frequency of spontaneously proliferated (SP) cells, identified by complete loss of CFSE, was significantly lower among Mll1KO donor cells than among WT donor cells (Figure 2C), indicating that MLL1 promotes efficient spontaneous proliferation under steady-state conditions. To exclude the possibility that the observed difference resulted from congenic markers rather than MLL1 deficiency, congenically marked WT (CD45.2^+^) and WT (CD45.1^+^) CD4^+^ T cells were co-transferred into WT recipients (Figure 2D). As expected, both donor populations generated comparable frequencies of SP cells (Figure 2E, F), confirming that the proliferative defect was specific to MLL1 deficiency.

**Figure 2.**
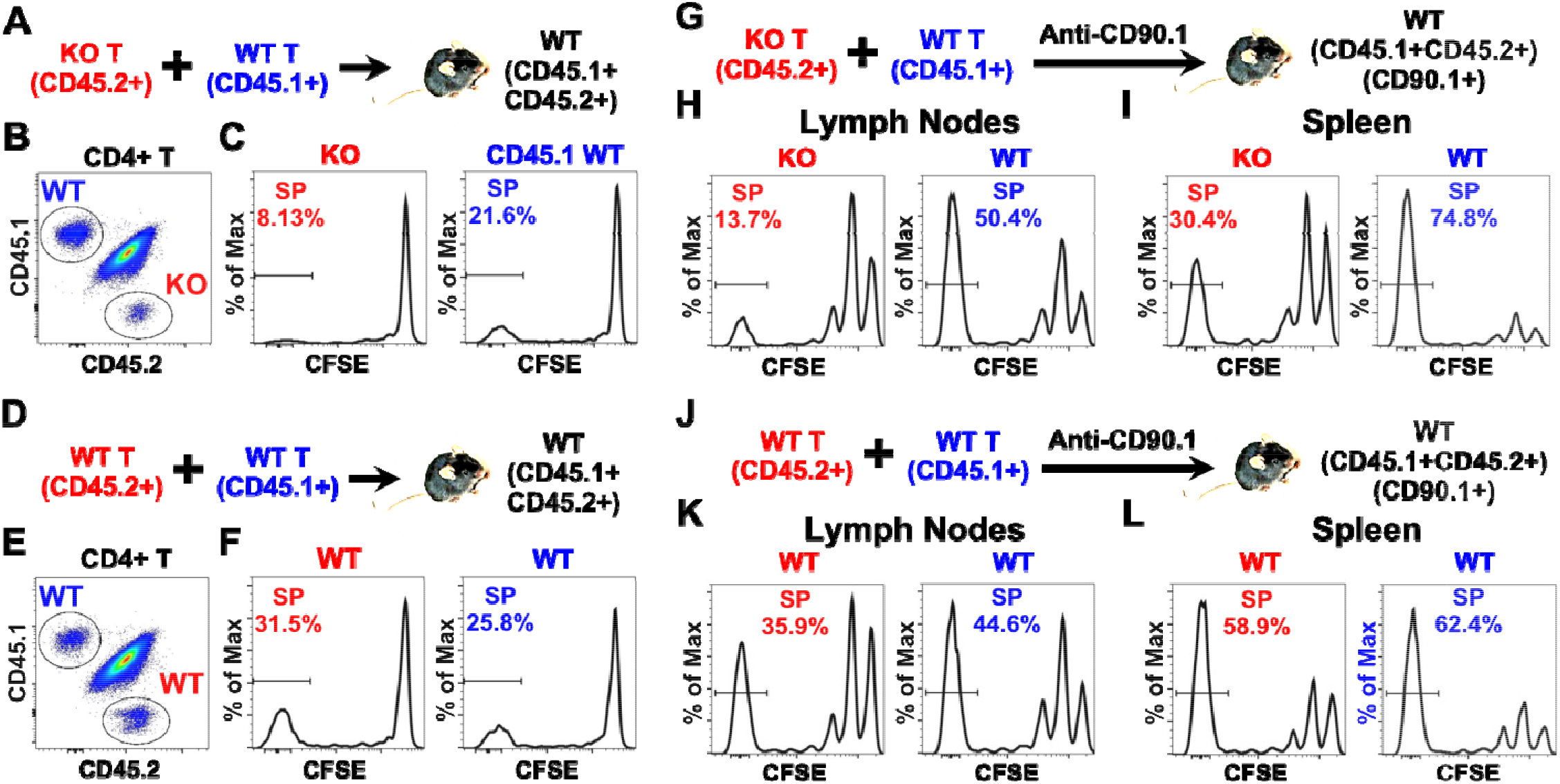
MLL1 promotes participation in spontaneous proliferation under competitive conditions. (A-C) Naïve T cells from congenically marked Mll1KO (CD45.2^+^) and wild-type (WT; CD45.1^+^) mice were mixed, labeled with CFSE, and transferred into unmanipulated WT (CD45.1^+^CD45.2^+^) recipients. Donor-derived cells were analyzed 5 weeks after transfer by flow cytometry. (A) Schematic of the experimental design. (B) Representative flow cytometric analysis of CD4^+^ T cells from peripheral lymph nodes showing CD45.1 and CD45.2 expression. (C) CFSE dilution in gated CD45.2^+^ Mll1KO and CD45.1^+^ WT CD4^+^ T cells. Spontaneously proliferated (SP) cells were defined as cells that had completely diluted CFSE. Data are representative of two independent experiments. (D-F) Control experiment using congenically marked WT (CD45.2^+^ and CD45.1^+^) T cells, analyzed as in (A-C). (D) Schematic of the experimental design. (E) Representative CD45.1/CD45.2 staining. (F) CFSE dilution in gated WT donor cells. Data are representative of two independent experiments. (G-I) CFSE-labeled Mll1KO (CD45.2^+^) and WT (CD45.1^+^) T cells were transferred into WT (CD45.1^+^CD45.2^+^CD90.1^+^) recipients, followed by anti-CD90.1 treatment. Donor cells were analyzed 3 weeks after transfer. (G) Schematic of the experimental design. (H, I) CFSE dilution in CD4^+^ T cells from peripheral lymph nodes (H) and spleen (I). Data are representative of two independent experiments. (J-L) Control experiment under lymphopenic conditions using congenically marked WT donor cells, analyzed as in (G-I). (J) Schematic of the experimental design. (K, L) CFSE dilution in CD4^+^ T cells from peripheral lymph nodes (K) and spleen (L). Data are representative of two independent experiments.

Spontaneous proliferation is constrained by competition for limiting resources and is enhanced under lymphopenic conditions [1, 11, 24]. We therefore asked whether reducing competition could rescue the proliferative defect of Mll1KO T cells. Congenically marked naïve Mll1KO (CD45.2^+^) and WT (CD45.1^+^) CD4^+^ T cells were co-transferred into WT (CD45.1^+^CD45.2^+^CD90.1^+^) recipients (Figure 2G). Recipient mice were treated with a depleting anti-CD90.1 antibody to selectively reduce endogenous T cells while sparing donor populations. Three weeks later, donor-derived cells were analyzed by flow cytometry. As expected, lymphopenia increased the frequency of SP cells in both donor populations compared with unmanipulated recipients. However, Mll1KO donor cells continued to generate significantly fewer SP cells than WT donor cells in both peripheral lymph nodes (Figure 2H) and spleen (Figure 2I). In contrast, no differences were observed between congenically marked WT donor populations subjected to the same treatment (Figure 2J-L). Thus, although reducing competition enhanced spontaneous proliferation overall, it failed to eliminate the disadvantage of Mll1KO T cells. Together, these findings demonstrate that MLL1 intrinsically promotes efficient spontaneous proliferation and provides a competitive advantage during the generation of MP CD4^+^ T cells under both steady-state and lymphopenic conditions.

### MLL1 maintains TCR expression during T-cell proliferation

Because spontaneous proliferation depends on TCR stimulation by self and environmental antigens [6], we asked whether MLL1 influences TCR expression during this process. We first compared surface TCR expression on naïve CD4^+^ T cells from Mll1KO and WT mice. As shown in Figure 3A, naïve Mll1KO and WT CD4^+^ T cells expressed comparable levels of TCR, indicating that MLL1 is not required for basal TCR expression in resting naïve T cells. We next examined whether TCR expression was maintained during spontaneous proliferation. Congenically marked naïve CD4^+^ T cells from Mll1KO (CD45.2^+^) and WT (CD45.1^+^) mice were labeled with CFSE and co-transferred into Rag1-deficient (Rag1KO) recipients (Figure 3B). Rag1KO mice were used because spontaneous proliferation occurs efficiently in lymphopenic hosts [7]. One week after transfer, a substantial fraction of donor cells had undergone spontaneous proliferation, as indicated by CFSE dilution (Figure 3C). Notably, WT CD4^+^ T cells maintained TCR expression following spontaneous proliferation, whereas Mll1KO CD4^+^ T cells exhibited a marked reduction in TCR expression (Figure 3C). Consequently, spontaneously proliferated (SP) Mll1KO CD4^+^ T cells expressed significantly lower levels of surface TCR than their WT counterparts (Figure 3D). In contrast, undivided naïve donor cells from both genotypes expressed comparable TCR levels.

**Figure 3.**
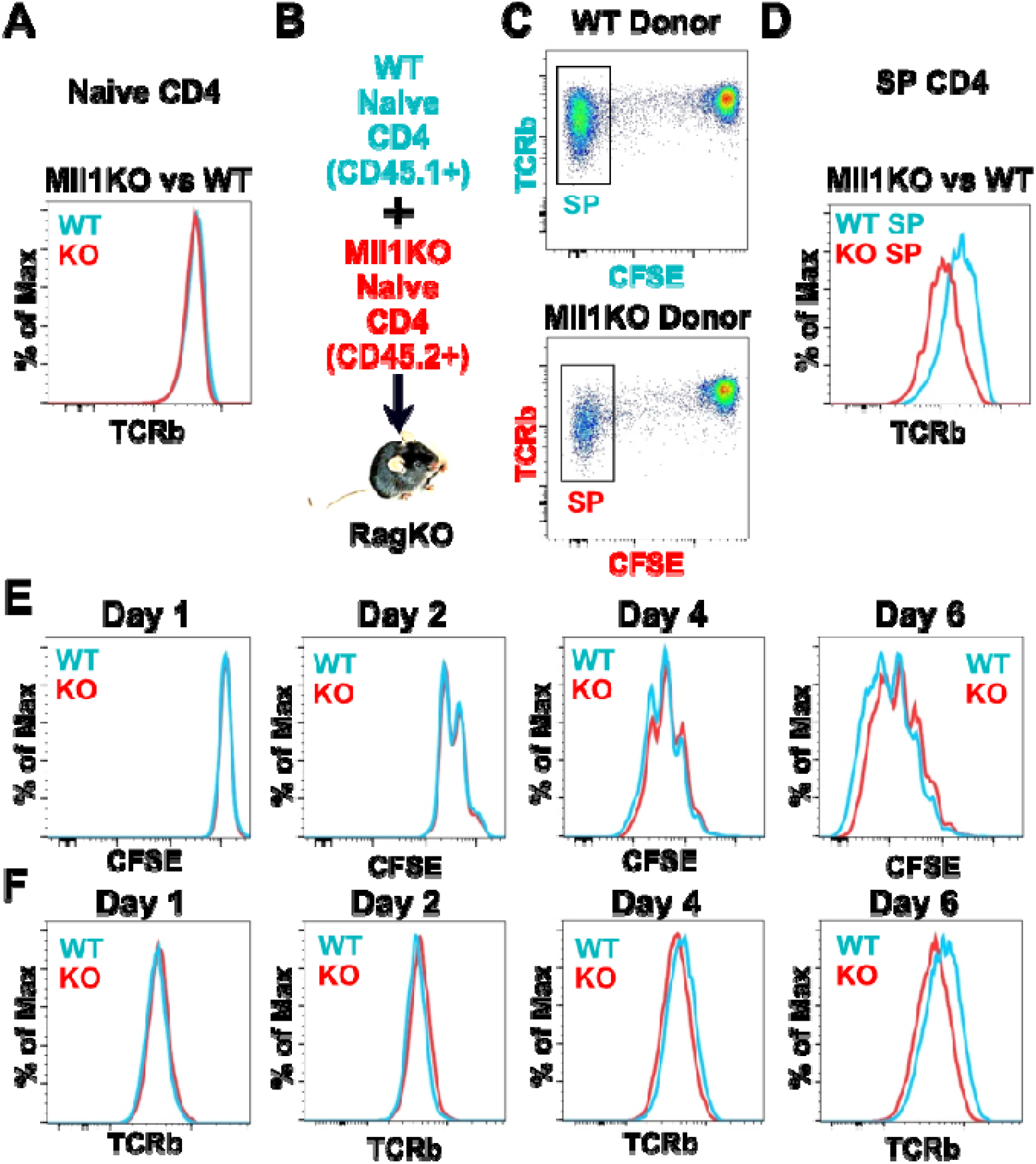
MLL1 maintains TCR expression during T-cell proliferation. (A) Naïve CD4^+^ T cells from Mll1KO mice and their WT littermates were analyzed by flow cytometry. The expression levels of TCR were compared. Data are representative of 4 independent experiments. (B-D) Congenically marked naïve CD4^+^ T cells were isolated from Mll1KO (CD45.2^+^) and WT (CD45.1^+^) mice, labeled with CFSE and co-transferred into Rag1KO recipients. Donor cells were analyzed 1 week after transfer by flow cytometry. (B) Experimental design. (C) Representative TCR expression on naïve (undivided) and spontaneously proliferated (SP) CD4^+^ T cells. (D) The expression levels of TCR on spontaneously proliferated (SP) WT and Mll1KO donor CD4^+^ T cells were compared. Data are representative of two independent experiments. (E-F) Naïve CD4^+^ T cells from Mll1KO and WT mice were labeled with CFSE and stimulated in vitro with anti-CD3ε and anti-CD28. Cells were analyzed by flow cytometry on days 1, 2, 4, and 6 after activation. (E) CFSE dilution as a measure of cell proliferation. (F) Surface TCR expression during activation. Data are representative of two independent experiments.

MLL1 functions as an epigenetic reader and mitotic bookmarking protein. The PHD3 domain of MLL1 binds H3K4me3 with high affinity and specificity [25], enabling MLL1 to associate with a large fraction of actively transcribed promoters [18, 19]. Through this mitotic bookmarking function, MLL1 facilitates the restoration of transcription following cell division [17]. We therefore asked whether the loss of TCR expression in Mll1KO cells was associated with cell proliferation itself rather than being unique to spontaneous proliferation. To address this question, CFSE-labeled naïve CD4^+^ T cells from Mll1KO and WT mice were stimulated in vitro with anti-CD3 and anti-CD28 and analyzed on days 1, 2, 4, and 6 after activation. Mll1KO and WT cells underwent comparable proliferation through day 4, with only a modest reduction in proliferation becoming apparent in Mll1KO cells by day 6 (Figure 3E). Surface TCR expression was also comparable between Mll1KO and WT cells during the early stages of activation (days 1 and 2; Figure 3F). However, by day 4, Mll1KO cells began to exhibit reduced TCR expression relative to WT cells, and this difference became more pronounced by day 6. Thus, loss of TCR expression in Mll1KO cells was observed not only during spontaneous proliferation in vivo but also during antigen receptor-driven proliferation in vitro. These findings indicate that MLL1 is required to maintain TCR expression during repeated rounds of cell division and are consistent with a role for MLL1-dependent transcriptional bookmarking in preserving TCR expression during proliferation.

Together, these findings demonstrate that MLL1 is dispensable for TCR expression in resting naïve CD4^+^ T cells but is required to maintain TCR expression during cell proliferation. The selective loss of TCR expression in dividing Mll1KO cells suggests that MLL1-dependent transcriptional maintenance becomes particularly important during repeated rounds of cell division, a defining feature of spontaneous proliferation and memory phenotype T-cell generation.

### Reduced TCR expression does not impair MP T-cell accumulation in the absence of WT competitors

The mixed bone marrow chimera experiments shown in Figure 1 demonstrated that Mll1KO CD4^+^ T cells were underrepresented within the MP compartment when they developed alongside WT T cells. To determine whether MLL1 is intrinsically required for MP T-cell generation and maintenance, we examined MP CD4^+^ T cells in unmanipulated Mll1KO mice. Unexpectedly, the frequency of memory phenotype (MP) CD4^+^ T cells was comparable between Mll1KO and WT mice in both peripheral lymph nodes (PLN; Figure 4A) and spleen (Figure 4D). Thus, in the absence of WT competitor cells, Mll1KO CD4^+^ T cells were capable of generating and maintaining a normal MP compartment. Despite normal MP-cell accumulation, Mll1KO MP CD4^+^ T cells expressed lower levels of surface TCR than WT MP cells in both PLN (Figure 4B) and spleen (Figure 4E). In contrast, naïve CD4^+^ T cells from Mll1KO and WT mice expressed comparable levels of TCR (Figure 4C, F). These findings indicate that MLL1 selectively maintains TCR expression in MP CD4^+^ T cells while being dispensable for TCR expression in naïve cells.

**Figure 4.**
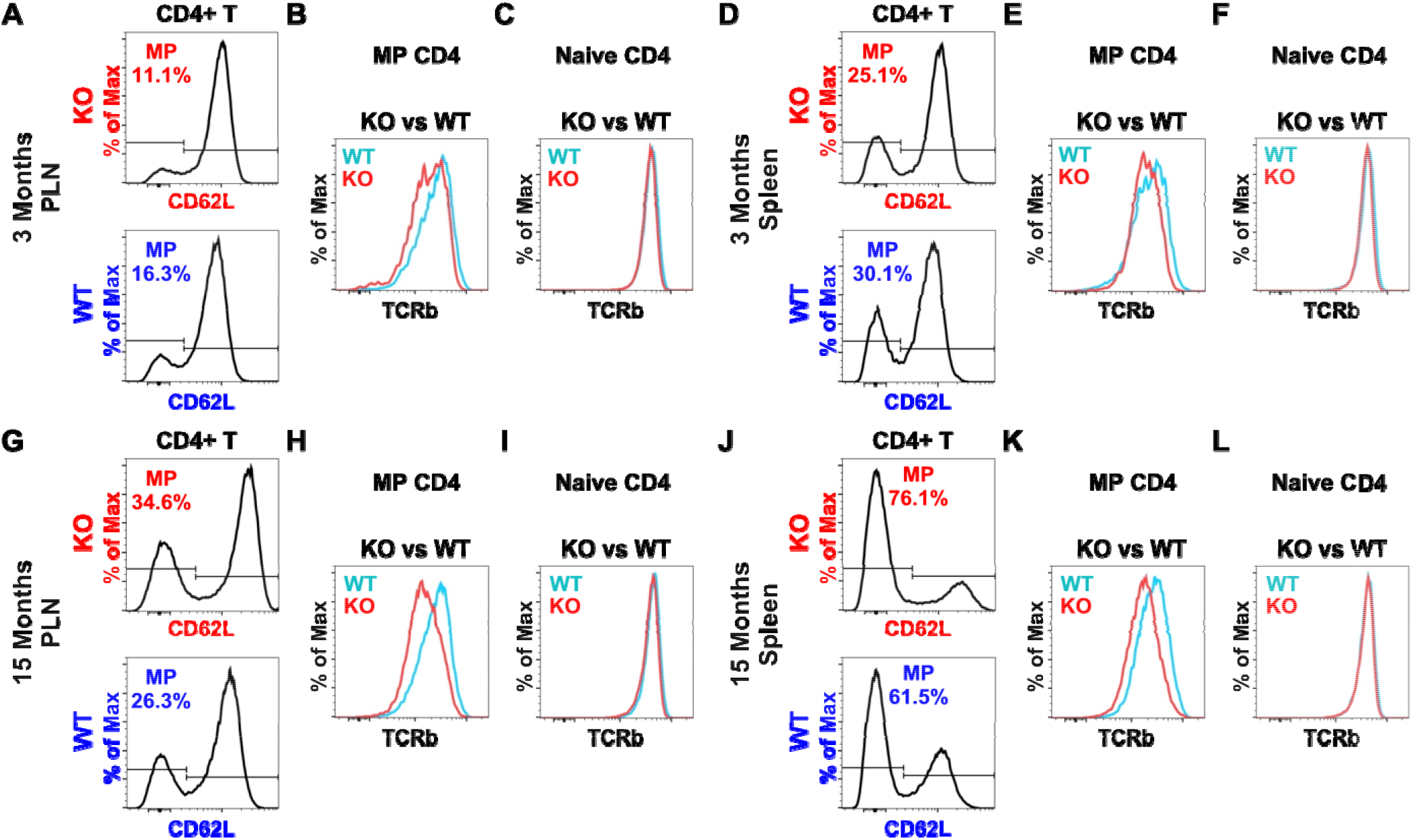
Reduced TCR expression does not impair MP T-cell accumulation in the absence of WT competitors. CD4^+^ T cells from peripheral lymph nodes (PLN) and spleen of 3-month-old (A-F) and 15-month-old (G-L) Mll1KO mice were compared with age-matched wild-type (WT) littermates by flow cytometry. (A, D, G, J) Frequency of CD62L^−^ MP cells within the CD4^+^ T cell population. (B, E, H, K) Surface TCR expression on MP (CD62L^−^) CD4^+^ T cells. (C, F, I, L) Surface TCR expression on naïve (CD62L^+^) CD4^+^ T cells. Data are representative of three independent experiments.

To determine whether MLL1 contributes to the age-associated expansion of the MP compartment, Mll1KO mice and WT littermates were aged to 15 months. As expected, WT mice exhibited a marked increase in MP-cell frequency with age in both PLN (Figure 4G compared with Figure 4A) and spleen (Figure 4J compared with Figure 4D). A similar age-dependent expansion was observed in Mll1KO mice, demonstrating that MLL1 is not intrinsically required for the accumulation of MP CD4^+^ T cells during aging. Although MP-cell frequencies were comparable between genotypes, reduced TCR expression persisted in aged Mll1KO MP T cells in both PLN (Figure 4H) and spleen (Figure 4K). In contrast, naïve CD4^+^ T cells from aged Mll1KO mice continued to express normal levels of TCR (Figure 4I and 4L). Thus, the selective requirement for MLL1 in maintaining TCR expression within the MP compartment is maintained throughout life.

Together, these findings demonstrate that MLL1 is not required for the generation or age-associated accumulation of MP CD4^+^ T cells in the absence of WT competitors. Rather, MLL1 selectively maintains TCR expression in MP T cells, suggesting that reduced TCR expression compromises competitive fitness rather than MP-cell development itself.

### MLL1 confers a competitive advantage during spontaneous proliferation

The observation that Mll1KO MP CD4^+^ T cells accumulated normally in unmanipulated mice (Figure 4) yet were underrepresented in mixed bone marrow chimeras (Figure 1), suggested that MLL1 might confer a competitive advantage during spontaneous proliferation rather than being absolutely required for the process. To determine whether Mll1KO T cells were intrinsically capable of undergoing spontaneous proliferation, CFSE-labeled naïve CD4^+^ T cells from Mll1KO and WT mice were transferred separately into Rag1-efficient (Rag1KO) recipients. Donor cells were recovered and analyzed one week later by flow cytometry. Strikingly, Mll1KO T cells underwent spontaneous proliferation to a similar extent as WT T cells (Figure 5A, B), demonstrating that MLL1 is not required for spontaneous proliferation in the absence of competing WT cells. However, despite proliferating efficiently, Mll1KO T cells failed to maintain normal levels of surface TCR expression following proliferation, whereas WT T cells retained high levels of TCR expression (Figure 5A, B). These findings indicate that impaired maintenance of TCR expression is a cell-intrinsic consequence of MLL1 deficiency.

**Figure 5.**
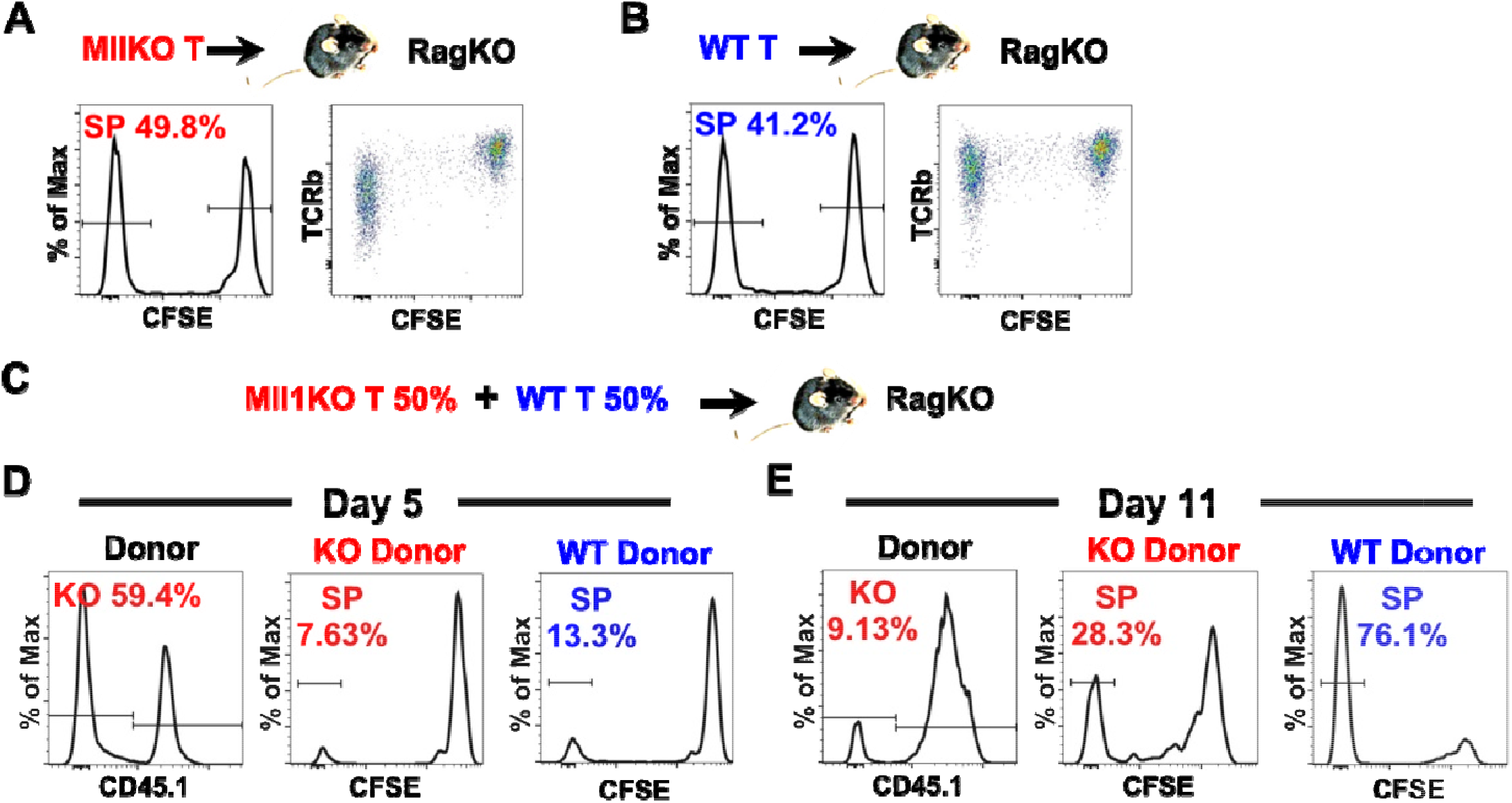
MLL1 confers a competitive advantage during spontaneous proliferation. (A-B) T cells from Mll1KO mice (A) or their WT littermates (B) were labeled with CFSE and transferred into Rag1KO recipients. One week later, donor cells were recovered and analyzed by flow cytometry. The percentage of spontaneously proliferated cells and the levels of TCR expression were determined. Data are representative of two independent experiments. (C-E) Congenically marked Mll1KO (CD45.2^+^) and wild-type (WT; CD45.1^+^) T cells were mixed at a 1:1 ratio, labeled with CFSE, and transferred into Rag1KO recipients. Five and 11 days later, donor cells were recovered and analyzed by flow cytometry. (C) Experimental design. (D-E) The representation of Mll1KO cells (KO) in the donor population and the percentage of spontaneously proliferated cells (SP) in Mll1KO and WT donor-derived T cells were determined at days 5 (D) and 11 (E) after transfer. Data are representative of three independent experiments.

We next asked whether reduced TCR expression affected the competitive fitness of Mll1KO T cells during spontaneous proliferation. Equal numbers of CFSE-labeled, congenically marked Mll1KO (CD45.2^+^) and WT (CD45.1^+^) naïve CD4^+^ T cells were mixed and transferred into Rag1KO recipients (Figure 5C . Donor cells were analyzed 5 and 11 days after transfer. Spontaneously proliferated cells first became detectable on day 5 (Figure 5D). At this early time point, Mll1KO and WT donor cells remained represented at approximately equal frequencies, and no significant differences in spontaneous proliferation were observed between the two populations. By day 11, however, the proportion of Mll1KO donor cells had declined markedly relative to WT donor cells (Figure 5E). This loss of representation was accompanied by a substantial reduction in the frequency of MP cells within the Mll1KO population.

Thus, although Mll1KO T cells are capable of undergoing spontaneous proliferation when transferred alone, they are progressively outcompeted by WT T cells when both populations coexist. Together, these findings demonstrate that MLL1 is not required for spontaneous proliferation itself but instead confers a competitive fitness advantage during this process.

### MLL1 sustains TCRα gene transcription during T-cell activation

To identify the transcriptional mechanism underlying the loss of TCR expression in proliferating Mll1KO T cells, we performed RNA-seq analysis on WT and Mll1KO CD4^+^ T cells four days after activation with anti-CD3 and anti-CD28. A total of 17,687 genes were analyzed, of which 1,053 were differentially expressed between WT and Mll1KO cells. Notably, a large fraction of the genes that were reduced in Mll1KO cells corresponded to Trav genes, which encode the variable regions of the TCRα chain (Figure 6). Multiple Trav family members exhibited significant decreases in expression, with many exhibiting substantial reductions in expression. In contrast, expression of other components of the TCR complex, including TCRβ and the CD3 signaling subunits (CD3δ, CD3γ, CD3ε, and CD3ζ), was largely unchanged.

**Figure 6.**
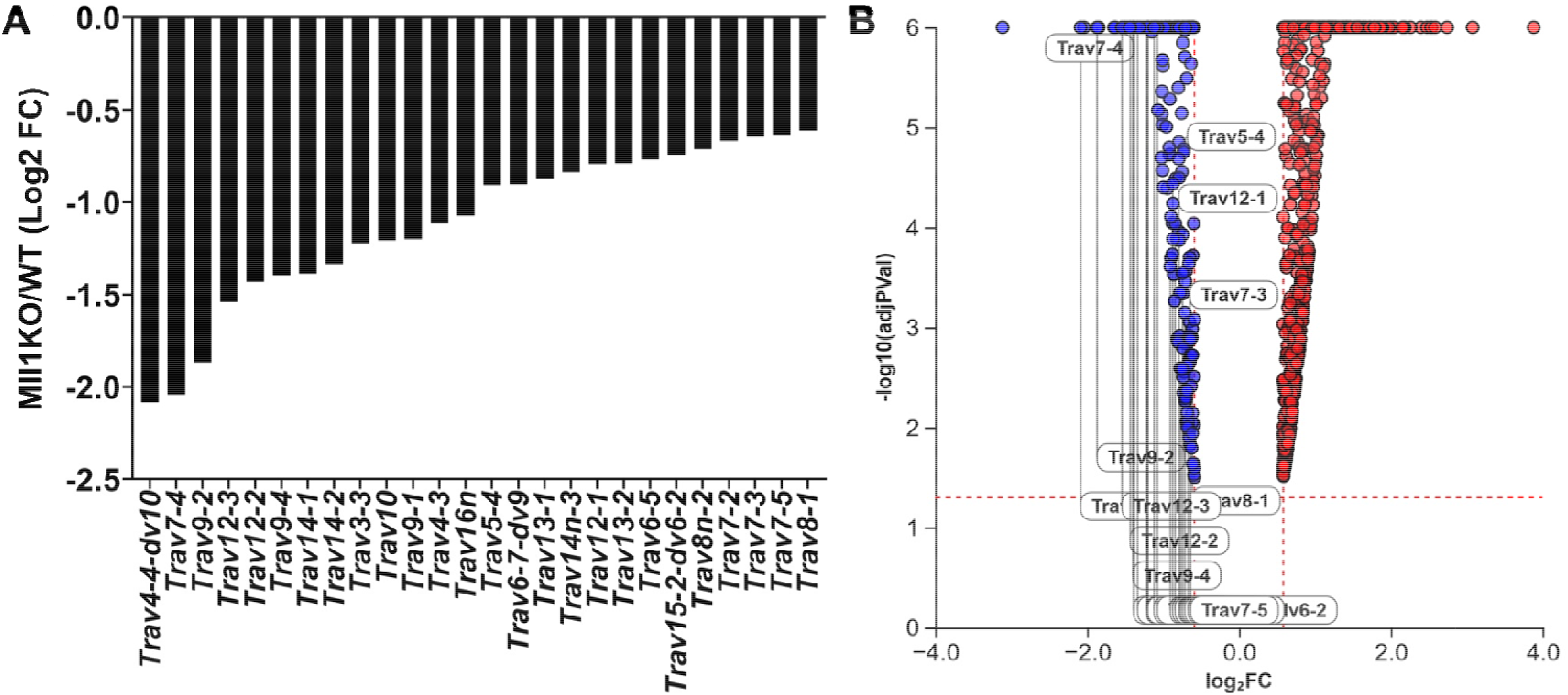
MLL1 sustains TCRα gene transcription during T-cell activation. Naïve T cells from T cell-specific Mll1KO mice and age-matched WT were activated in vitro with anti-CD3e and anti-CD28. On day 4 of the activation, cells were collected and subjected to RNA-Seq analysis. (A) Bar graph showing the base-2 logarithm of the fold change (Log2FC). (B) Volcano plot showing the statistical significance of the changes.

These findings demonstrate that MLL1 is required to maintain transcription of a broad array of TCRα variable region genes during T-cell activation. The selective reduction of Trav transcription provides a molecular explanation for the progressive loss of surface TCR expression observed in proliferating Mll1KO T cells and supports a model in which MLL1 preserves T-cell competitive fitness by sustaining TCRα gene expression during repeated rounds of cell division.

### MLL1 preserves competitive fitness of MP Treg cells

The CD4^+^ T cell compartment consists of regulatory T (Treg) cells, defined by expression of the transcription factor Foxp3, and conventional CD4^+^ T cells, which lack Foxp3 expression. To determine the role of MLL1 in Treg cells, we first analyzed Treg cells in unmanipulated Mll1KO mice. As shown in Figure 7A, the frequency of Treg cells within the CD4 T cell population was comparable between Mll1KO and WT mice. Similarly, the proportion of memory phenotype (MP) Treg cells within the Treg cell population was comparable between Mll1KO and WT mice (Figure 7B). However, surface TCR expression was reduced on Mll1KO MP Treg cells relative to WT controls (Figure 7C), whereas naïve Treg cells showed no difference in TCR expression (Figure 7D). These findings indicate that, as in conventional CD4^+^ T cells, MLL1 is specifically required to maintain TCR expression in MP but not naïve Treg cells. In the absence of WT competitors, a moderate reduction TCR expression does not impair MP Treg cell accumulation.

**Figure 7.**
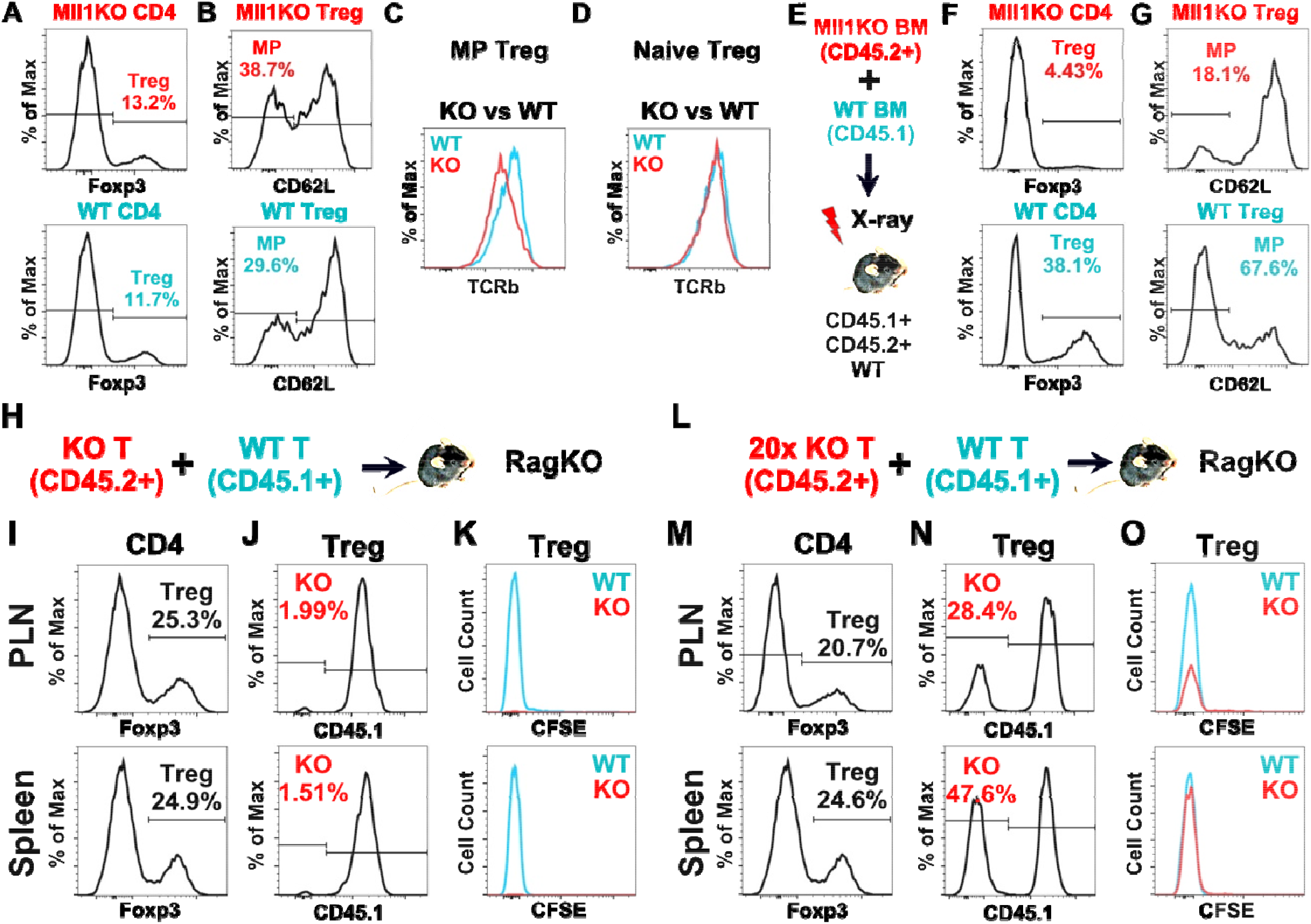
MLL1 promotes competitive fitness of MP Treg cells. (A-D) CD4^+^ T cells from the spleen of T cell-specific Mll1KO mice were compared with age-matched wild-type (WT) littermates by flow cytometry. (A) Frequency of Treg cells within the CD4^+^ T cell population. (B) Frequency of memory phenotype (MP; CD62L^−^) cells within the Treg cell population. (C) Surface TCR expression on MP Treg cells. (D) Surface TCR expression on naïve (CD62L^+^) Treg cells. Data are representative of three independent experiments. (E-G) Bone marrow cells (BM) from congenically marked Mll1KO (CD45.2^+^) and wild-type (WT; CD45.1^+^) mice were mixed and transferred into lethally irradiated WT (CD45.1^+^CD45.2^+^) recipients. Donor-derived T cells were analyzed by flow cytometry 4 months later. (E) Experimental design. (F) Frequency of Treg cells within the CD4^+^ T cell population. (G) Frequency of memory phenotype (MP; CD62L^−^) cells within the Treg cell population. (H-K) Congenically marked Mll1KO (CD45.2^+^) and wild-type (WT; CD45.1^+^) T cells were mixed at a 1:1 ratio, labeled with CFSE, and transferred into Rag1KO recipients. Four weeks later, donor cells were recovered and analyzed by flow cytometry. (H) Experimental design. The percentage of Treg cells within the donor CD4 population (I), the percentage of Mll1KO donor-derived cells within the Treg cell population (J) and the CFSE dilution profile of Mll1KO donor-derived and WT donor-derived Treg cells (K) in the spleen and peripheral lymph nodes (PLN) were determined. Data are representative of 2 independent experiments. (L-O) Congenically marked Mll1KO (CD45.2^+^) and wild-type (WT; CD45.1^+^) T cells were mixed at a 20:1 ratio, labeled with CFSE, and transferred into Rag1KO recipients. Four weeks later, donor cells were recovered and analyzed as in (H-K). (L) Experimental design. The percentage of Treg cells within the donor CD4 population (M), the percentage of Mll1KO donor-derived cells within the Treg cell population (N) and the CFSE dilution profile of Mll1KO donor-derived and WT donor-derived Treg cells (O) in the spleen and peripheral lymph nodes (PLN) were determined. Data are representative of 2 independent experiments.

To determine the role of MLL1 in MP Treg cells under competitive conditions, we generated mixed bone marrow chimeras using bone marrow from congenically marked Mll1KO (CD45.2+) and WT (CD45.1+) mice. Lethally irradiated CD45.1+CD45.2+ WT mice were used as BM recipients (Figure 7E). Donor Treg cells were analyzed 4 months after bone marrow transfer. As shown in Figure 7F, the percentage of Treg cells within Mll1KO CD4+ T cells was much lower than that in WT CD4+ T cells (4.43% vs 38.1%). Furthermore, the percentage of MP cells within the Mll1KO Treg cell population was much lower than that in the WT Treg cell population (18.1% vs 67.6%) (Figure 7G). These data indicate that MLL1 confers a competitive advantage that promotes accumulation of MP Treg cells.

To determine whether spontaneous proliferation of Treg cells is similarly regulated by MLL1, equal numbers of congenically marked Mll1KO (CD45.2^+^) and WT (CD45.1^+^) T cells were labeled with CFSE and transferred into Rag1KO recipients (Figure 7H). Four weeks later, donor-derived T cells were recovered and analyzed by flow cytometry. Treg cells were readily detected in both peripheral lymph nodes and spleen (Figure 7I). Remarkably, WT cells accounted for nearly the entire Treg compartment, whereas Mll1KO cells represented less than 2% of donor-derived Treg cells (Figure 7J). Nearly all WT Treg cells had completely diluted CFSE, indicating extensive spontaneous proliferation (Figure 7K). Because too few Mll1KO Treg cells were recovered for meaningful analysis, it remained unclear whether Mll1KO Treg cells failed to proliferate or were simply outcompeted by WT cells.

To distinguish between these possibilities, the experiment was repeated using a 20-fold excess of Mll1KO donor cells relative to WT cells (Figure 7L). Increasing the input ratio substantially increased the representation of Mll1KO cells within the recovered Treg compartment (Figure 7N) without altering the overall frequency of Treg cells among donor-derived CD4^+^ T cells (Figure 7M). Under these conditions, nearly all Mll1KO Treg cells completely diluted CFSE, similar to WT Treg cells (Figure 7O), indicating that Mll1KO Treg cells retain the capacity to undergo spontaneous proliferation.

Together, these findings demonstrate that MLL1 is not required for spontaneous proliferation of Treg cells per se. Rather, MLL1 confers a substantial competitive fitness advantage that enables MP Treg cells to accumulate in the presence of WT competitors.

### Homeostasis is lost when MLL1 is missing in Treg cells but not in conventional T cells

The preceding experiments demonstrated that MLL1 maintains the competitive fitness of both conventional and regulatory CD4^+^ T cells by preserving TCR expression during proliferation. Because immune homeostasis depends on a balance between autoreactive conventional CD4^+^ T cells and Treg cells [16], we hypothesized that disruption of this balance would occur when MLL1 deficiency was restricted to Treg cells but not conventional T cells.

To test this possibility, we generated Treg-specific MLL1-deficient mice (Mll1KO(Treg)) by crossing Mll1fl/fl mice [20] with Foxp3YFP-Cre mice [26]. In contrast to T cell-specific MLL1-deficient mice (Mll1KO(T)), which remained healthy throughout life, Mll1KO(Treg) mice failed to thrive and became moribund by approximately 6 months of age (Figure 8A). Mll1KO(Treg) mice also developed pronounced lymphadenopathy and splenomegaly (Figure 8B). Histological examination revealed extensive leukocytic infiltration of peripheral tissues, including the lung, whereas no comparable pathology was observed in Mll1KO(T) mice (Figure 8C). These findings demonstrate that selective loss of MLL1 in Treg cells is sufficient to disrupt immune homeostasis and cause severe inflammatory disease. In contrast, simultaneous loss of MLL1 in both Treg and conventional T cells is well tolerated. Thus, the pathological consequence of MLL1 deficiency depends not simply on the absolute function of Treg cells but on the relative balance between MLL1-deficient Treg cells and MLL1-sufficient conventional T cells.

**Figure 8.**
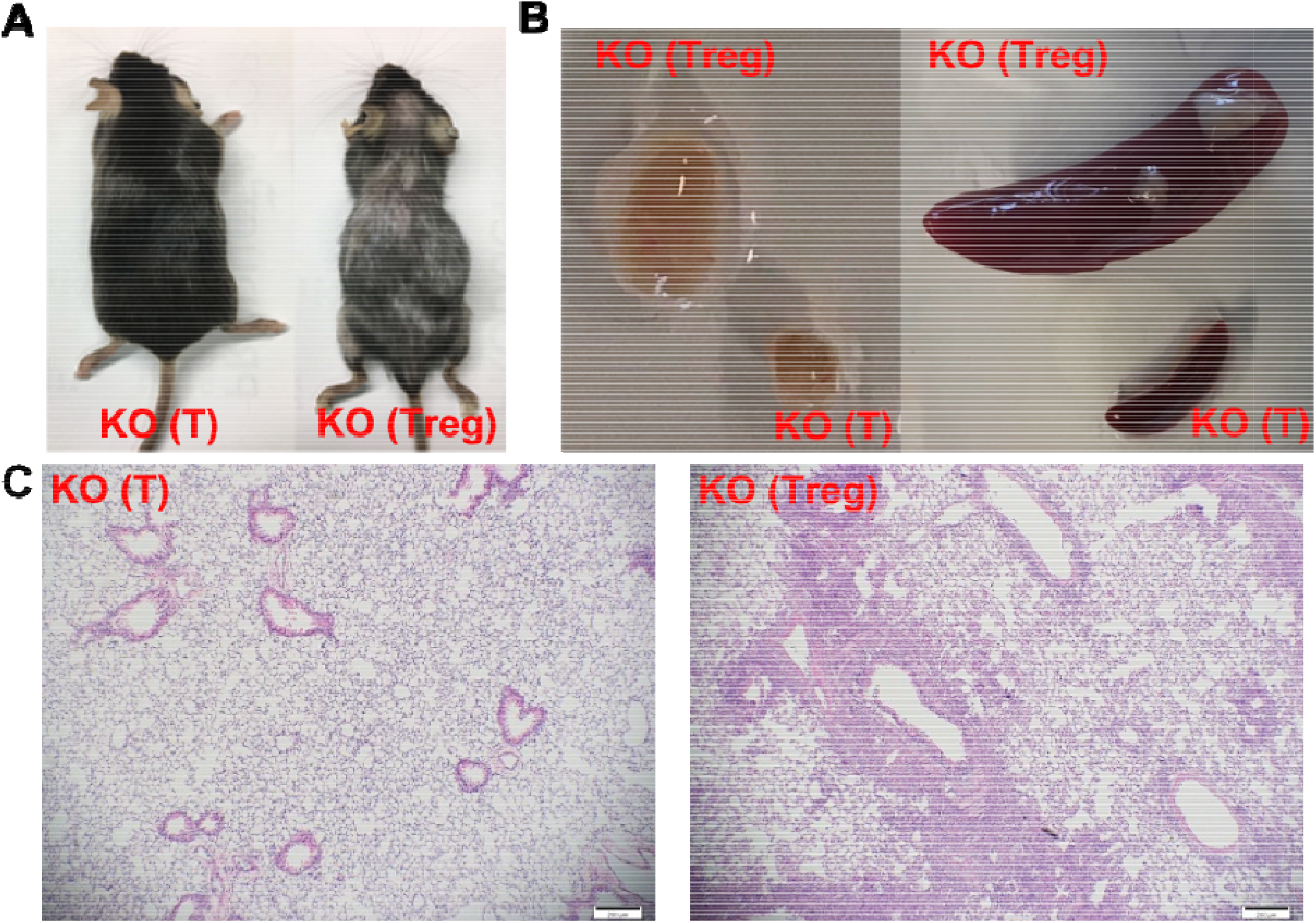
Homeostasis is lost when MLL1 is missing in Treg cells but not in conventional T cells. Four-month-old T cell-specific MLL1-deficient mice (KO (T)) were compared with age-matched Treg-specific MLL1-deficient mice (KO (Treg)). (A) Gross physical appearance. (B) Size of peripheral lymph nodes and spleen. (C) Representative histological sections of lung tissue. Data are representative of more than 10 paired mice.

Together with the competitive fitness defects described above, these results support a model in which MLL1 maintains immune tolerance by preserving the ability of MP Treg cells to compete effectively with autoreactive conventional CD4^+^ T cells.

### Memory phenotype Treg cells restore immune homeostasis during aging

To determine whether WT Treg cells could compensate for the loss of Mll1KO(Treg) cells and restore immune homeostasis, we generated mixed bone marrow chimeras using congenically marked Mll1KO(Treg) (CD45.2^+^) and WT (CD45.1^+^) donor cells. Lethally irradiated CD90.1^+^ WT mice were used as recipients. Because preliminary experiments indicated that Mll1KO(Treg)-derived Treg cells declined over time, donor bone marrow was mixed at a ratio of 70% Mll1KO(Treg) and 30% WT cells prior to transfer (Figure 9A). Unlike unmanipulated Mll1KO(Treg) mice, which developed lethal inflammatory disease with age, all mixed chimeras remained healthy, indicating that WT donor-derived cells restored immune homeostasis. Twelve months after bone marrow transfer, the frequency of Treg cells remained within the normal range (Figure 9B). As expected, WT-derived conventional CD4^+^ T cells represented approximately 20% of the total conventional CD4^+^ T-cell compartment, consistent with their representation in the donor bone marrow inoculum (Figure 9C). In contrast, WT-derived cells accounted for more than 90% of the Treg compartment (Figure 9D). Notably, the majority of WT-derived Treg cells exhibited a memory phenotype (CD62L^−^), whereas approximately half of the residual Mll1KO(Treg)-derived Treg cells retained a naïve phenotype (Figure 9E). These findings indicate that restoration of immune homeostasis was associated with preferential expansion of WT memory phenotype Treg cells.

**Figure 9.**
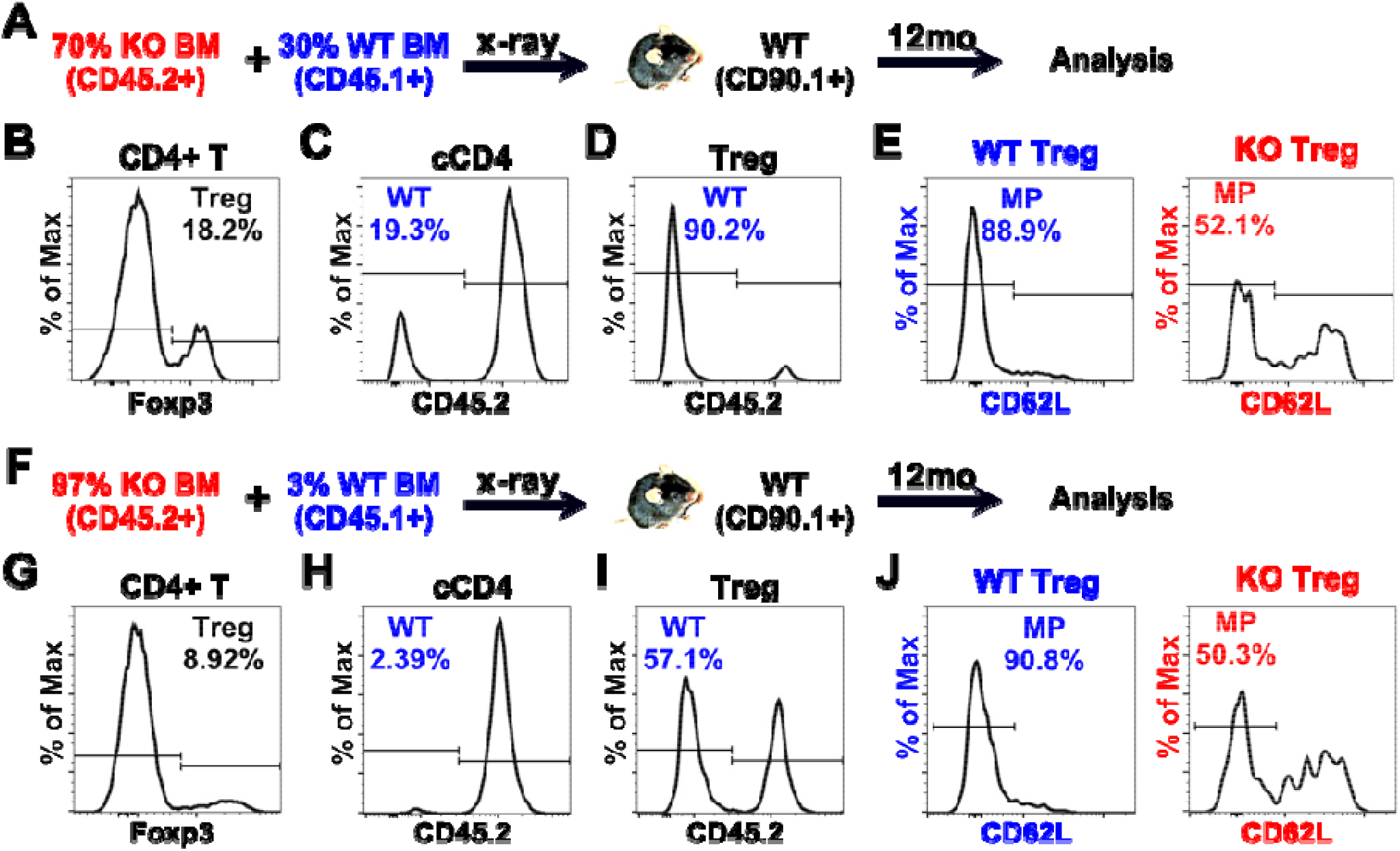
Memory phenotype Treg cells restore immune homeostasis during aging. (A-E) Bone marrow cells from congenically marked Treg-specific Mll1KO(Treg) (CD45.2^+^) and WT (CD45.1^+^) mice were mixed at a ratio of 70:30 and transferred to lethally irradiated WT (CD90.1^+^) hosts. Twelve months later, donor-derived T cells were recovered and analyzed by flow cytometry. After gating out the residual host (CD90.1^+^) cells, Mll1KO (KO) and WT cells were identified based on their expression of CD45.2. (A) Experimental design. (B) Treg cells and conventional CD4 (cCD4) T cells were identified in the CD4+ T cells population based on their expression of Foxp3. (C) Percentage of WT-derived cells within the conventional CD4^+^ T cell population. (D) Percentage of WT-derived cells within the Treg cell population. (E) Percentage of memory phenotype (MP) cells within the WT-derived and Mll1KO(Treg)-derived Treg cell population. Data are representative of 2 experiments. (F-J) Bone marrow cells from congenically marked Treg-specific Mll1KO (CD45.2^+^) and WT (CD45.1^+^) mice were mixed at a ratio of 97:3 and transferred to lethally irradiated WT (CD90.1^+^) hosts. Twelve months later, donor-derived T cells were recovered and analyzed by flow cytometry as in (B-E). (F) Experimental design. (G) Treg cells and conventional CD4 (cCD4) T cells were identified in the CD4+ T cells population based on their expression of Foxp3. (H) Percentage of WT-derived cells within the conventional CD4+ T cell population. (I) Percentage of WT-derived cells within the Treg cell population. (J) Percentage of memory phenotype (MP) cells within the WT-derived and Mll1KO(Treg)-derived Treg cell population. Data are representative of 2 experiments.

To determine the extent to which WT Treg cells could compensate for Mll1KO(Treg) deficiency, mixed chimeras were generated using donor bone marrow containing only 3% WT cells and 97% Mll1KO(Treg) cells (Figure 9F). Ten chimeras were generated in two independent experiments. Eight developed lethal inflammatory disease, whereas two survived and aged normally. Analysis of the surviving mice 12 months after reconstitution revealed a normal frequency of Treg cells within the CD4^+^ T-cell compartment (Figure 9G). WT-derived conventional CD4^+^ T cells represented approximately 3% of the conventional T-cell compartment, consistent with their contribution to the donor inoculum (Figure 9H). Remarkably, however, WT-derived cells constituted more than half of the Treg compartment (Figure 9I). As observed in the 70:30 chimeras, this selective expansion was driven predominantly by memory phenotype Treg cells (Figure 9J).

Together, these findings demonstrate that memory phenotype Treg cells possess a remarkable capacity for expansion and can restore immune homeostasis even when derived from a very small pool of WT precursors. These results identify expansion of the memory phenotype Treg-cell compartment as a critical mechanism for maintaining immune tolerance during aging.

## Discussion

Immune tolerance must be maintained throughout life despite profound age-associated changes in the T-cell compartment. One of the most prominent of these changes is thymic involution, which progressively reduces the production of naïve T cells. Nevertheless, the peripheral CD4^+^ T-cell compartment remains remarkably stable because memory phenotype (MP) T cells accumulate with age [11-13]. Although this phenomenon has been recognized for many years, the biological significance of MP T-cell expansion has remained unclear. In the present study, we identify competitive expansion of MP Treg cells as a critical mechanism for maintaining immune tolerance during aging. Using MLL1 deficiency as a model, we show that modest reductions in T-cell competitive fitness impair the accumulation of MP Treg cells and disrupt immune homeostasis. These findings reveal an unexpected link between epigenetic maintenance of TCR expression, intra-clonal competition, and age-associated preservation of immune tolerance.

Previous studies have established that MP CD4^+^ T cells arise through spontaneous proliferation driven by self and environmental antigens [7, 8]. Because spontaneous proliferation depends on TCR signaling, MP-cell development is constrained by competition among T cells for limiting stimulatory signals. Our findings demonstrate that this competition is not simply a mechanism regulating the size of the MP compartment but instead plays an important role in determining which cells persist within it. Mll1-deficient T cells remained capable of spontaneous proliferation and generated MP populations normally when transferred in the absence of WT competitors. However, when WT and Mll1KO cells coexisted, Mll1KO cells were progressively lost from the MP compartment. These observations indicate that MLL1 is not required for spontaneous proliferation itself but rather for competitive fitness during spontaneous proliferation. The distinction between proliferative capacity and competitive fitness is central to understanding the phenotype observed in this study. Mll1KO conventional CD4^+^ T cells and Treg cells retained the capacity to undergo spontaneous proliferation when competitor cells were absent. The defect became apparent only under competitive conditions. Thus, maintenance of MP populations is governed not simply by the ability to proliferate but by the ability to compete successfully for antigen-dependent signals over time.

A major finding of this study is that relatively modest reductions in TCR expression can profoundly influence competitive fitness. MLL1 deficiency selectively reduced TCR expression in MP cells while leaving naïve T cells largely unaffected. This reduction was observed during spontaneous proliferation in vivo and during activation-induced proliferation in vitro. Mechanistically, MLL1 maintained transcription of multiple Trav genes encoding the variable regions of the TCRα chain. In contrast, expression of TCRβ and CD3 signaling components was largely preserved. These findings identify maintenance of TCRα transcription as a previously unrecognized MLL1-dependent function in proliferating T cells. Because spontaneous proliferation is driven by continual antigen recognition, even modest reductions in TCR expression are expected to reduce the efficiency with which cells compete for limiting antigenic stimuli. Over time, small differences in competitive fitness are amplified, resulting in progressive loss of Mll1KO cells from the MP compartment.

Our findings further suggest that intra-clonal competition serves analogous functions in the thymus and the periphery. During thymic development, competition among T cells with similar antigen specificities contributes to the establishment of the balance between Treg cells and autoreactive conventional CD4^+^ T cells [9, 10]. The present study suggests that this balance must continue to be maintained throughout life. In the peripheral MP compartment, Treg cells and autoreactive conventional CD4^+^ T cells appear to remain subject to antigen-dependent competition. Under these conditions, preservation of competitive fitness becomes essential for maintaining the equilibrium between regulatory and pathogenic populations. Thus, intra-clonal competition may not simply shape the T-cell repertoire during development but may continuously regulate immune homeostasis throughout life.

The consequences of impaired competitive fitness were particularly striking in Treg cells. In unmanipulated Mll1KO mice, Treg-cell frequencies remained normal despite reduced TCR expression. However, under competitive conditions, Mll1KO Treg cells were profoundly disadvantaged and failed to generate a normal MP Treg compartment. Selective deletion of MLL1 in Treg cells resulted in severe inflammatory disease, whereas simultaneous deletion of MLL1 in both conventional and regulatory T cells was well tolerated. These findings indicate that immune tolerance depends not on the absolute fitness of either population but on the balance between them. When both populations experience a comparable reduction in competitive fitness, homeostasis is maintained. In contrast, when only Treg cells are disadvantaged, autoreactive conventional T cells gain a relative advantage, resulting in loss of tolerance and inflammatory disease.

The experiments involving mixed bone marrow chimeras provided additional insight into the role of MP Treg cells during aging. Remarkably, a relatively small number of WT Treg precursors was sufficient to restore immune homeostasis in mice dominated by Mll1-deficient Treg cells. In surviving chimeras, WT Treg cells expanded far beyond their initial representation within the donor inoculum and ultimately constituted the majority of the Treg compartment. This expansion was mediated predominantly by MP Treg cells. These findings demonstrate that MP Treg cells possess a remarkable capacity for self-renewal and expansion and suggest that this population serves as an important reservoir for maintaining immune tolerance as thymic production of naïve Treg cells declines.

Age-associated accumulation of MP Treg cells is frequently viewed as a contributor to immune dysfunction in older individuals because excessive Treg activity can suppress responses to infection, vaccination, and cancer [15]. While this interpretation is likely correct, our findings suggest that expansion of MP Treg cells also serves an essential physiological function. As thymic involution progressively reduces the supply of naïve Treg cells, maintenance of tolerance becomes increasingly dependent on expansion of existing MP Treg cells. Thus, age-associated accumulation of MP Treg cells may represent an adaptive response that preserves self-tolerance in the face of declining thymic function. The beneficial and detrimental consequences of MP Treg-cell expansion may therefore represent two sides of the same biological process.

In summary, our findings identify competitive expansion of MP Treg cells as a critical mechanism for maintaining immune tolerance during aging. We show that preservation of TCR expression is required for competitive fitness during spontaneous proliferation and that MLL1 supports this process by maintaining transcription of TCRα genes during cell division. More broadly, these results suggest that intra-clonal competition continues to regulate the balance between regulatory and autoreactive T cells throughout life and becomes increasingly important as thymic output declines. Thus, competitive expansion of MP Treg cells may represent a fundamental mechanism by which immune tolerance is preserved during aging.

## Materials and Methods

### Mice

*Mll1*^*flox/flox*^ mice have been described previously [20]. CD4-Cre transgenic mice and *Foxp3*^*YFP-Cre*^ transgenic mice (B6.129(Cg)-*Foxp3*^*tm4(YFP/icre)Ayr*^/J) were purchased from the Jackson Laboratory (Bar Harbor, ME) [26]. All strains were on the C57BL/6 (CD45.2+CD45.1-) mouse background. T cell-specific MLL-deficient mice (Mll1KO) were generated by crossing the *Mll1*^*flox/flox*^ mice with the CD4-Cre mice. Treg-specific MLL1 knockout mice (Mll1KO (Treg)) by crossing *Mll1*^*flox/flox*^ mice with the *Foxp3*^*YFP-Cre*^ transgenic mice. Male CD45.1+ C57BL/6 congenic mice (B6.SJL-*Ptprc*^*a*^ *Pepc*^*b*^/BoyJ), Rag1 deficient (RagKO) mice (B6.129S7-*Rag1*^*tm1Mom*^/J) and D90.1+ C57BL/6 congenic mice (B6.PL-*Thy1*^*a*^/Cy) were purchased from the Jackson Laboratory. C57BL/6 F1 congenic mice (CD45.1+CD45.2+) were produced by crossing male CD45.1+ C57BL/6 congenic with female C57BL/6 (CD45.1-CD45.2+) mice. Mice were individually identified by ear tags and maintained under specific pathogen-free conditions and provided food and water ad libitum. The University of Michigan Committee on Use and Care of Animals (UCUCA) approved all animal studies.

### Generation of mixed bone marrow chimeras

Mixed bone marrow chimeras were generated as previously described [27]. Congenically marked recipient mice were irradiated with a single dose of 9 Gy. Lineage-depleted bone marrow cells (CD4^−^, CD8^−^, CD5^−^, CD19^−^, B220^−^, NK1.1^−^, and CD11b^−^) from congenically marked donor mice were mixed at the indicated ratios, assessed for viability, and transferred intravenously within 2 h of irradiation.

### Flow cytometry

Flow cytometric analyses were performed as previously described [27]. Briefly, single-cell suspensions were prepared from spleens, lymph nodes, and peripheral blood. Spleens and lymph nodes were mechanically dissociated, whereas peripheral blood samples were subjected to red blood cell lysis prior to staining.

Approximately 1 × 10L cells were stained with fluorochrome-conjugated antibodies in PBS containing 5% BSA. Cells were washed in PBS containing 1% BSA and fixed/permeabilized using the Foxp3 Transcription Factor Fixation/Permeabilization kit (Thermo Fisher Scientific) according to the manufacturer’s instructions. Intracellular staining was performed in PBS containing 5% BSA. Following staining, cells were washed and resuspended in PBS containing 1% BSA before acquisition on a BD LSR II flow cytometer. Data were analyzed using FlowJo software (Tree Star).

Antibodies used included anti-CD19 (6D5), anti-CD62L (MEL-14), anti-CD44 (IM7), anti-CXCR3 (CXCR3-173), anti-TCRβ (H57-597), anti-CD49d (9C10), anti-CD122 (TM-b1), anti-CD5 (53-7.3), anti-CD6 (J90-462), anti-CD69 (H1.2F3), anti-CD127 (A7R34), anti-H-2KL (28-8-6), anti-Ki-67 (SolA15), anti-CD4 (GK1.5), anti-CD11b (M1/70), anti-CD45 (30-F11), anti-CD45.1 (A20), anti-CD45.2 (104), and anti-CD8β (H35-17.2). Antibodies were obtained from BioLegend, eBioscience, Bio-Rad (formerly Serotec), or Cell Signaling Technology.

### T cell adoptive transfer

Adoptive transfer experiments were performed as previously described [28]. Briefly, single-cell suspensions were prepared from spleens and lymph nodes. Unwanted cell populations were depleted using biotinylated antibodies and anti-biotin magnetic negative-selection kits (Miltenyi Biotec, Auburn, CA). In some experiments, cells were labeled with CFSE prior to transfer. Viable cells were transferred intravenously in 0.5 ml PBS. For long-term tracking experiments, at least 5 × 10L viable cells were transferred per recipient. In selected experiments, recipient mice were treated intraperitoneally with 0.5 mg anti-CD90.1 antibody (clone 19E12) to transiently deplete endogenous T cells.

### In vitro T-cell activation

In vitro T-cell activation was performed as previously described [29]. Briefly, CD4^+^ T cells were isolated from spleens and peripheral lymph nodes and labeled with CFSE. Cells were stimulated using Dynabeads Mouse T-Activator CD3/CD28 (Thermo Fisher Scientific) in complete medium supplemented with IL-2 (20 ng/ml).

Dynabeads were added at a bead-to-cell ratio of 5:1. Samples were collected at the indicated time points and analyzed by flow cytometry.

### RNA sequencing

Total RNA was isolated using RNeasy Plus Micro kits (Qiagen) according to the manufacturer’s instructions. Residual genomic DNA was removed using RNase-Free DNase (Qiagen). RNA samples were submitted to the University of Michigan Advanced Genomics Core for library preparation, sequencing, and bioinformatic analysis. Differential gene expression analysis was performed using DESeq2 [30], using a negative binomial generalized linear model (thresholds: linear fold change >1.5 or <-1.5, Benjamini-Hochberg FDR (Padj) <0.05).

### Statistical analysis

Statistical analyses were performed using Prism software (GraphPad). Comparisons between two groups were performed using the unpaired Student’s t test. P values < 0.05 were considered statistically significant.

## Notes

### Competing Interest Statement

The authors have declared no competing interest.

